# Protozoan predation selects for key symbiotic traits in an environmentally transmitted beneficial symbiosis

**DOI:** 10.64898/2026.02.25.707974

**Authors:** Daravuth Cheam, Eshaine Sun, Isabel Jones, Isabella Ma, Miranda Magdaleno, Michele K. Nishiguchi

## Abstract

Beneficial associations between bobtail squids (Cephalopoda: Sepiolidae) and *Vibrio* bacteria encompass a unique association where symbionts are obtained environmentally from the surrounding environment. *Vibrio* symbionts are susceptible to a number of ecological pressures such as protozoan grazing whilst in their free-living state. Impacts of grazing have several consequences for symbiosis characteristics such as biofilm formation, a trait crucial for survival both in and outside the squid. Therefore, in order to ascertain how biotic factors such as grazing in the environment effect symbiotic success, two *V. fischeri* strains, ES114 and ETBB1-C were experimentally evolved in separate biofilm grazing experiments with the amoeba, *Acanthamoeba castellanii* and ciliate *Tetrahymena pyriformis*. Both ES114 and ETBB1-C biofilms were evolved up to 50 generations through serial passaging. At 50 generations, ES114 biofilms displayed variability in response to predation by both predators, whereas ETBB1-C biofilms were more stable across generations of grazing. *A. castellanii* decreased in population numbers when co-inoculated with ETBB1-C, whereas T. pyriformis increased in numbers with biofilm growth. Growth of *V. fischeri* biofilms in the presence of grazers such as *T. pyriformis* has an important role in inducing biofilm growth by acting as a chaperone for recycling nutrients back into the environment. Additionally, *V. fischeri* colonization fitness in the host was dependent on which grazer was used to evolve the biofilms. Such variation in response by *V. fischeri* to different types of predation demonstrates the versatility of this symbiont in its free living state and has subsequent impacts on the eventual association with squids.

**Importance:** This manuscript demonstrates the importance of biotic factors (such as protozoan grazing) in the environment that effect host colonization in a beneficial symbiosis. Using an experimental evolution approach, this work demonstrates how symbiotic biofilms can adapt to pressures such as grazing that subsequently influences the ability to colonize its invertebrate host.

## Introduction

Symbiotic interactions are crucial to life on Earth since microbes are found in association with all living organisms. Beneficial associations otherwise known as mutualisms in particular are interesting due to the nature of how both host and microbe benefit from one another (1, 2). Many beneficial associations play a significant role in virtually all living organisms, having some type of interaction with their host that benefits both partners (3). This is especially important given that many organisms depend on processes that are only driven by microbes in the environment but in turn, provide a necessary function for the host organism. For example, the human gut microflora is a major site where beneficial microbes are found and function in breaking down and digesting afood for their hosts (4). Studying the mechanisms that comprise the interaction between environmentally transmitted microbes and their partners is of particular importance given how the Earth’s climate is rapidly changing and effecting large ecosystems dependent upon microbial function.

A number of model organisms have been developed to study environmentally transmitted symbioses, particularly ones where the host and microbe are found both separately and in association with one another. One such system is the mutualism between sepiolid squids (Cephalopoda: Sepiolidae) and their bioluminescent symbiotic bacteria in the genus *Vibrio* (g-Proteobacteria: Vibrionaceae) (5). Host squids provide shelter and nutrients that allow them to grow four times faster than in the environment (6). In return, *Vibrio* bacteria provide bioluminescence that the squid uses in a behavior called counterillumination (7). This allows the squid to match downwelling moonlight from above to hide its silhouette from predators below as the squid searches for food and evade predators at night.

In the sepiolid squid-*Vibrio* mutualism, symbiotic bacteria must form biofilm-like aggregates on the inner surface of the squid light organ in order to colonize the light organ (8). Once the aggregates are pulled into the pores of the naïve juvenile light organ, planktonic *Vibrio* cells migrate and ultimately reside in the crypt spaces of the squid light organ (9). *Vibrio* bacteria then produce biofilm in the crypt spaces in order to persist inside the light organ (10). During the night, *Vibrio* bacteria grow four times faster than when in their free-living state in seawater (6) till they completely fill the light organ. Later at dawn, 95% of these bacteria are then vented out with the onset of light, and the remaining 5% persisting in the light organ to recolonize for the next diel cycle. Once outside the host, these vented bacteria can also form biofilms when colonizing a substrate and are susceptible to a number of ecological factors. One such biotic factor is protozoan predation (Fig. 1). Protozoan predators are known to have a dramatic effect on bacterioplankton populations, and can significantly change bacterial population structure and resilience in the free-living state. They are also impacted by abiotic factors that are associated with climate change (11), thereby having an indirect impact on symbiotic systems such as those associated with *Vibrio* bacteria.

**Figure 1.**
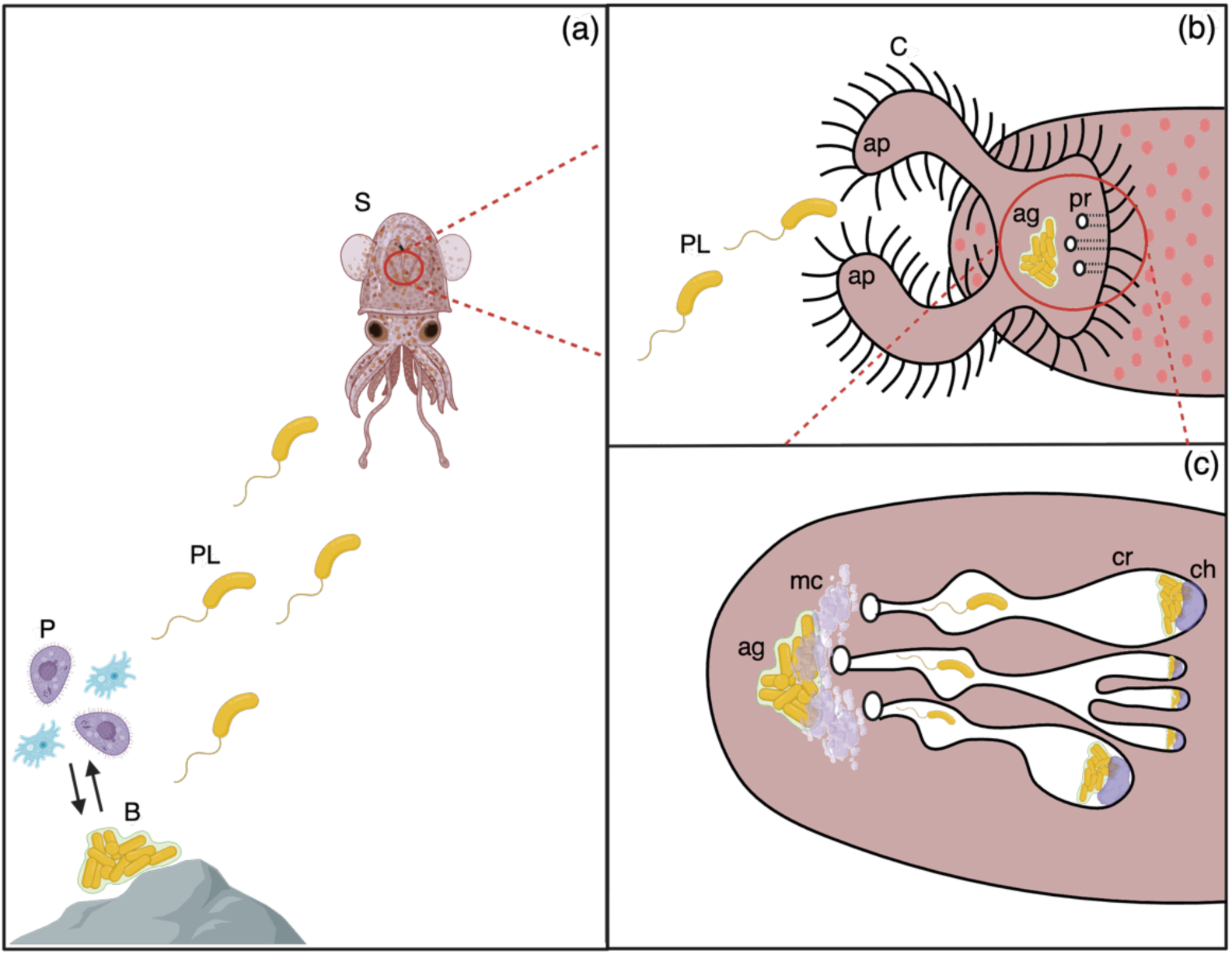
Illustration of *V. fischeri* life stages from free-living biofilms to host-associated symbionts. (a) *V. fischeri* biofilms, B, form outside of the squid host on substrates such as rocks. The biofilms are susceptible to ecological factors such as grazing by protozoans, P. When biofilms grow into their mature stage, they can release planktonic cells, PL, which can colonize their squid host, S. (b) The planktonic cells are brought towards the pore, pr, by appendages, ap, covered by cilia, C. When close to the pores, they are able to form biofilm-like aggregates, ag. (c) The aggregates form by attaching to a mucus layer, mc, around the pores. Aggregates in their mature form release planktonic cells that can go through the pores making their way to the crypts, cr, where they can proliferate around chitin food sources, ch.

Previous studies have examined protozoan grazing on other species of *Vibrio*, including *V. cholerae* and *V. fischeri* (12, 13). These studies demonstrated mechanisms of preventing protozoan attack by producing chemicals that ward off or kill their grazers. For example, *V. fischeri* biofilms responded differently to protozoan grazing depending on whether they were symbiotic or free-living strains. These studies found that biofilms of free-living strains were able to resist grazing by ciliates compared to symbiotic-competent biofilms, which were susceptible to grazers (13). Another study found that the closely related congener *V. cholerae* became more competent in colonization when passaged through expelled food vacuoles of protozoans (14). Although protozoans have not been reported to infect the light organ of bobtail squids, cephalopods in general are known to be infected by amoeba-like and ciliated protozoans (15). This study demonstrated that protozoans were reported to sometimes co-infect with Gram-negative bacteria. Since *V. fischeri* have a biphasic life history strategy where they are subject to host selection inside the squid while facing abiotic pressures outside the squid, balancing the fitness costs of both host and environment is crucial for *Vibrio* symbionts to persist over time (13). Whether evolutionary success in one stage provides an advantage to the other phase of the life cycle has not been closely examined, yet can provide a better perspective on beneficial associations that are transmitted through the environment. Therefore, we applied an experimental evolutionary approach to examine how free-living *V. fischeri* biofilms under continuous protozoan predation effects their ability to colonize, infect, and persist in their host squids. We measured physiological features such as growth rate, bioluminescence, motility, and host colonization which are key indicators of a successful symbioses.

## Methods

### Bacterial and protozoan strains

Bacterial strains and protozoan predators used in study are listed in Table 1. Bacterial strains were isolated from squids collected from either *Euprymna scolopes* (Hawaii-ES114) or *E. tasmanica* (Australia-ETBB1C). Protozoans Acanthamoeba castellanii and Tetrahymena pyriformis were obtained from the American Type Culture Collection (ATCC). *V. fischeri* were streaked from frozen glycerol stock cultures onto seawater tryptone (SWT) agar plates (5 g/L tryptone, 3 g/L yeast extract, 0.375% glycerol, 21 g/L Instant Ocean©, 7 g/L marine mix, and 15 g/L agar in water) to obtain isolated colonies. Plates were then incubated at 28°C for 24 hours. Protozoans *Acanthamoeba castellanii* and *Tetrahymena pyriformis* were kept in stock cultures made from peptone yeast glucose (5 g/L proteose peptone, 0.25 g/L yeast extract, and 5% D-glucose in 10% M9 minimal media).

**Table 1.** (a) Bacterial strains used in this study, their host organism, and habitat. (b) Protozoan predators used in this study are listed here, alongside with their mode of grazing, and where their ATCC product number.

| ***V. fischeri* strain** | **Squid Host** | **Location** |
| --- | --- | --- |
| ES114 | *Euprymna scolopes* | USA (Kaneohe Bay, O'ahu, HI) |
| ETBB1-C | *Euprymna tasmanica* | Australia (Botany Bay, Sydney, NSW) |

(b)
| **Organism** | **Grazer-type** | **ATCC Product Number** |
| --- | --- | --- |
| *Acanthamoeba castellanii* | Surface-feeder | 30010 |
| *Tetrahymena pyriformis* | Suspension-feeder | 30202 |

### Growth rate assay

Biofilms growth rates were determined through *in vitro* growth assays. These were completed by growing and collecting *V. fischeri* biofilms of each strain in triplicate and measured every 4 hours up to 24 hours. All samples were rinsed with 34 ppt seawater prior to collection to remove planktonic cells. After rinsing, the biofilms were resuspended in 500 µL of seawater and collected. They were then vortexed with a sterile glass bead to further resuspend the biofilm. Afterwards, they were diluted and plated on SWT agar plates. The plates were incubated for 24 hours at 28°C to allow for colonies to grow. Colonies were counted after each timepoint using a Stuart Colony Counter SC6+. For example, the biofilms in triplicate were harvested and quantified after 8 hours of incubation for the 8 hour timepoint. A growth curve was then generated using the Growthcurver package from R, from which the growth rate was calculated by fitting the growth curve data to a logistic equation (16).

### Grazing assays

Biofilm grazing assays are outlined in the diagram shown in Figure 2. Both *V. fischeri* ES114 and ETBB1C were individually grown and infected with either species of protozoans. In these grazing assays, *V. fischeri* ES114 or ETBB1C were first grown in SWT broth (same composition as SWT agar but without agar) overnight. Strains were then sub-cultured the next morning into fresh SWT media. Subcultures were grown until they reached their growth phase at approximately OD_600_ = 0.3. This corresponds to roughly 3 x 10^7^ CFU/mL assuming 1 OD is equivalent to 1 x 10^8^ CFU/mL (6) and represents the concentration of the initial biofilm inoculum. 500 mL aliquots of the subcultures were inoculated into 24-well plates and incubated at 28°C for 24 hours to allow biofilms to form. Six lineages of each treatment of both non-grazed and grazed biofilms were created. Six biological replicates were then used per lineage throughout the grazing assay.

**Figure 2.**
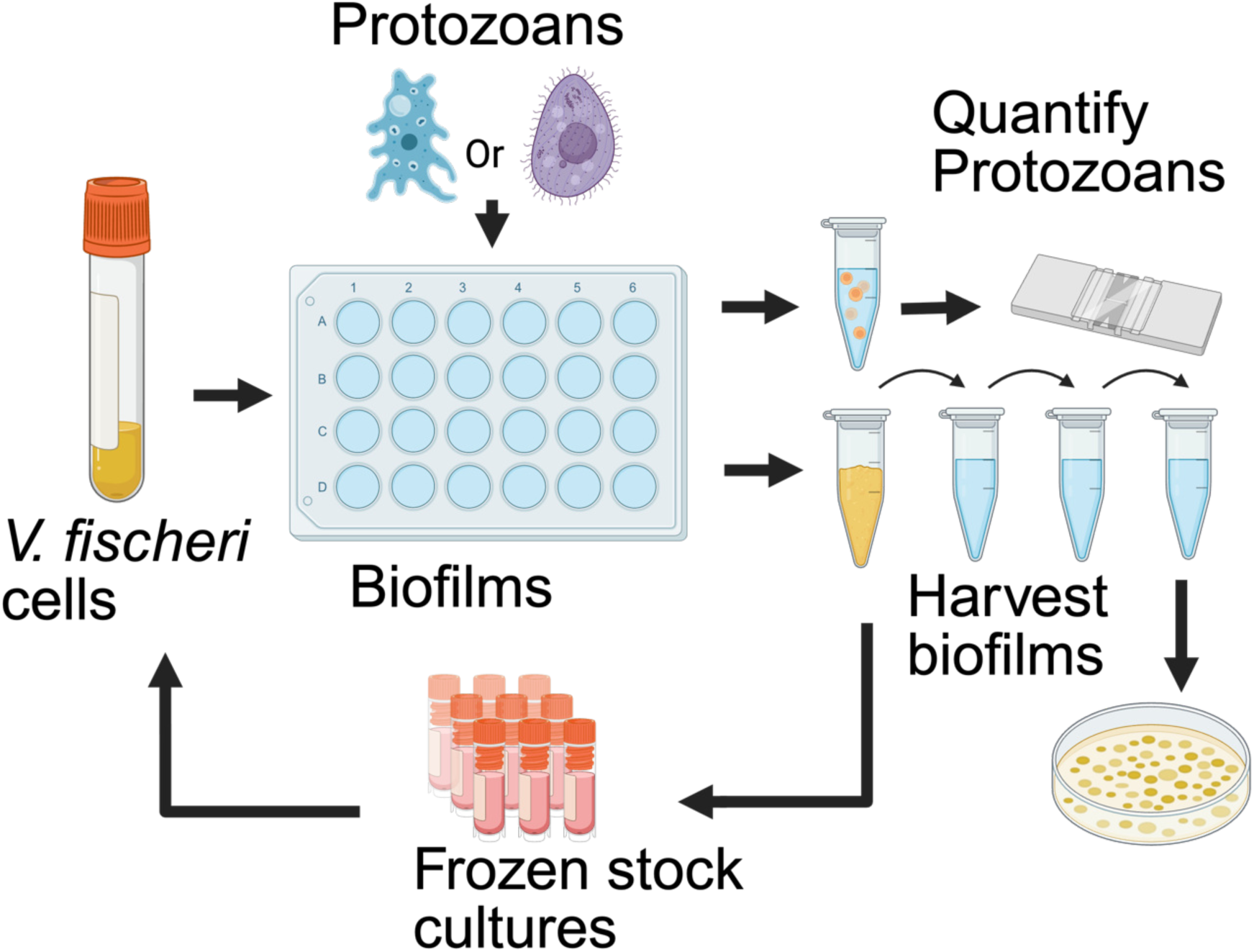
Experimental design of the biofilm grazing assay. *Vibrio fischeri* are grown in SWT liquid culture, which were then used as the inoculum to grow biofilms in 24-well plates. After biofilms have grown, they were treated with two types of protozoan predators (grazed biofilms). Another set of biofilms were untreated to serve as negative controls (non-grazed biofilms). Grazed and non-grazed biofilms were incubated for 24 hours. Afterwards, both protozoans and biofilms were harvested. Protozoans were quantified by hemocytometry, and biofilms were diluted and quantified by the colony count method.

After biofilm formation, the planktonic cells were removed and replaced with either seawater to serve as a negative control (non-grazed) biofilms, or with one of the two predators as the treatment group for grazed biofilms. Predators were added onto biofilms with an inoculum of 1,000 cells/mL. Nongrazed and grazed biofilms were then incubated for another 24 hours at 28°C without shaking. After 24 hours, the planktonic cells of non-grazed biofilms were removed and rinsed with 34 ppt seawater. Wells were then replaced with fresh seawater and subsequently collected. Planktonic cells for *T. pyriformis* grazed biofilms were also collected as *T. pyriformis* are generally found in suspension. The planktonic cells in *A. castellanii* grazed biofilms however were removed as *A. castellanii* are generally more associated with the biofilms and not the planktonic solution. All grazed treatments for biofilms were rinsed with seawater. Biofilms were then resuspended in 500 mL of 34 ppt seawater and subsequently collected. The strains are notated in the following manner: L = Lineage, Gn = generation, AC = *A. castellanii*, TP = *T. pyriformis*, NG = Nongrazed, and G = Grazed.

### Quantification of biofilms and protozoans

Biofilms were resuspended in 34 ppt seawater and then diluted 10,000-fold and plated on SWT agar plates and incubated overnight at 28°C to allow colonies to grow. The colonies were then enumerated as colony forming units (CFUs) using a colony counter. Protozoans were counted via hemocytometry after each grazing treatment. The remaining culture of resuspended biofilms were frozen in a final concentration of 20% glycerol for long-term storage at -80°C as fossil records and to compare against future grazing treatments.

### Bioluminescence Assays

Bioluminescence was measured by growing biofilms in white opaque 24-well plates for 24 hours at 28°C. The remaining planktonic cells were removed and replaced with 34 ppt sterile seawater. The 24-well plates were then placed in a SpectraMax iD5 plate reader which measured bioluminescence for each culture.

### Competition Experiments

*V. fischeri* strains were grown in SWT broth overnight. The cultures were then subcultured and allowed to grow until they reached an OD_600_ = 0.5. The two competing strains (ancestor vs. evolved) were then diluted in 34 ppt seawater. This inoculum (2500 CFUs of each strain/mL) of *V. fischeri* was aliquoted into 10 mL glass scintillation vials. Individual newly hatched squid were then placed into each vial. The inoculation mixture was replaced with fresh seawater after 12 hours of incubation. Bioluminescence was measured using a TD-20/20 Luminometer at 24 and 48 hours and seawater was replaced prior to each measurement. At 48 hours, hatchlings were euthanized in 2% ethanol (17), homogenized and plated onto SWT agar plates whereby colony counts were obtained from infected light organs (18). A competition index (1) was used for analysis between the two competing strains (19).

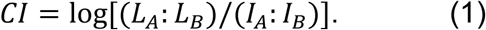

According to this equation, *CI* is the competition index, *L_A_*is the concentration of one strain and *L_B_* is the concentration of the other strain being competed with post-infection. *I_A_*and *I_B_* are the concentrations of both strains in the initial inoculum prior to infection.

### Statistical Analysis

Graphical representation and statistical analysis of the grazing assays, bioluminescence assays, and competition experiments were performed using GraphPad Prism 10. The level of significance was based on P-values which are categorized by asterisks. Smaller P-values represents greater significance. If a comparison is not significant, then “ns” for not significant is used.

## Results

Growth rates of the initial cultures of *V. fischeri* ES114 and ETBB1-C biofilms were found to be 0.375 gen/hr and 0.417 gen/hr, respectively. These growth rates were converted to gen/day since grazing assays were executed over the span of 24 hours. Therefore, growth rates of ES114 and ETBB1-C biofilms are equivalent to 9 gen/day and 10 gen/day respectively. From these growth rates, an expected evolutionary trajectory of both biofilms strains can estimate number of generations (Figs. 3, 4).

**Figure 3.**
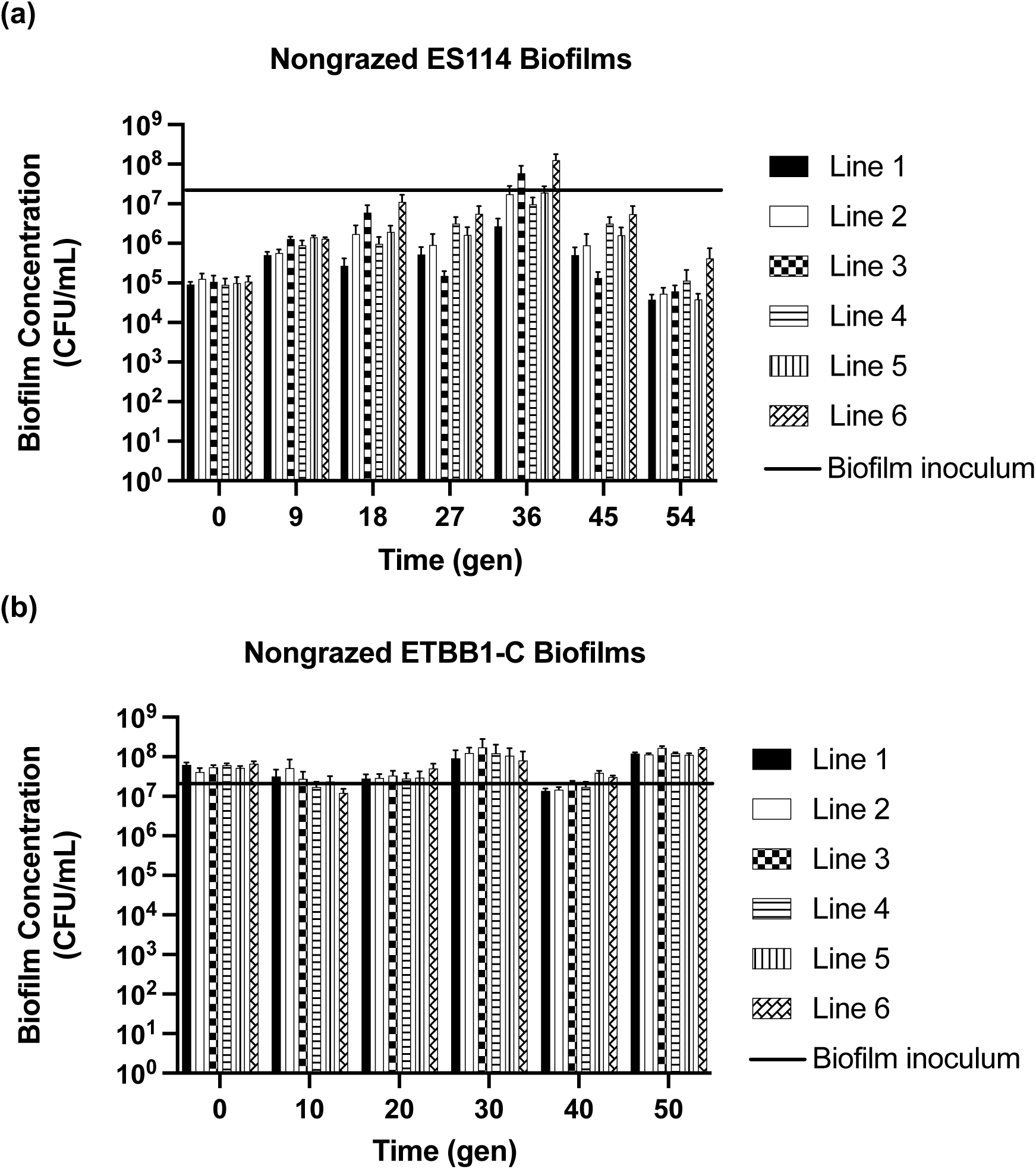
Concentrations of nongrazed a) ES114, and b) ETBB1-C biofilms across generations for six independent lineages. The initial biofilm concentration of around 3 x 10^7^ CFU/mL is represented by the black bar. Error bars are based on the standard error of the mean (SEM). The population of each lineage is composed of six biological replicates.

**Figure 4.**
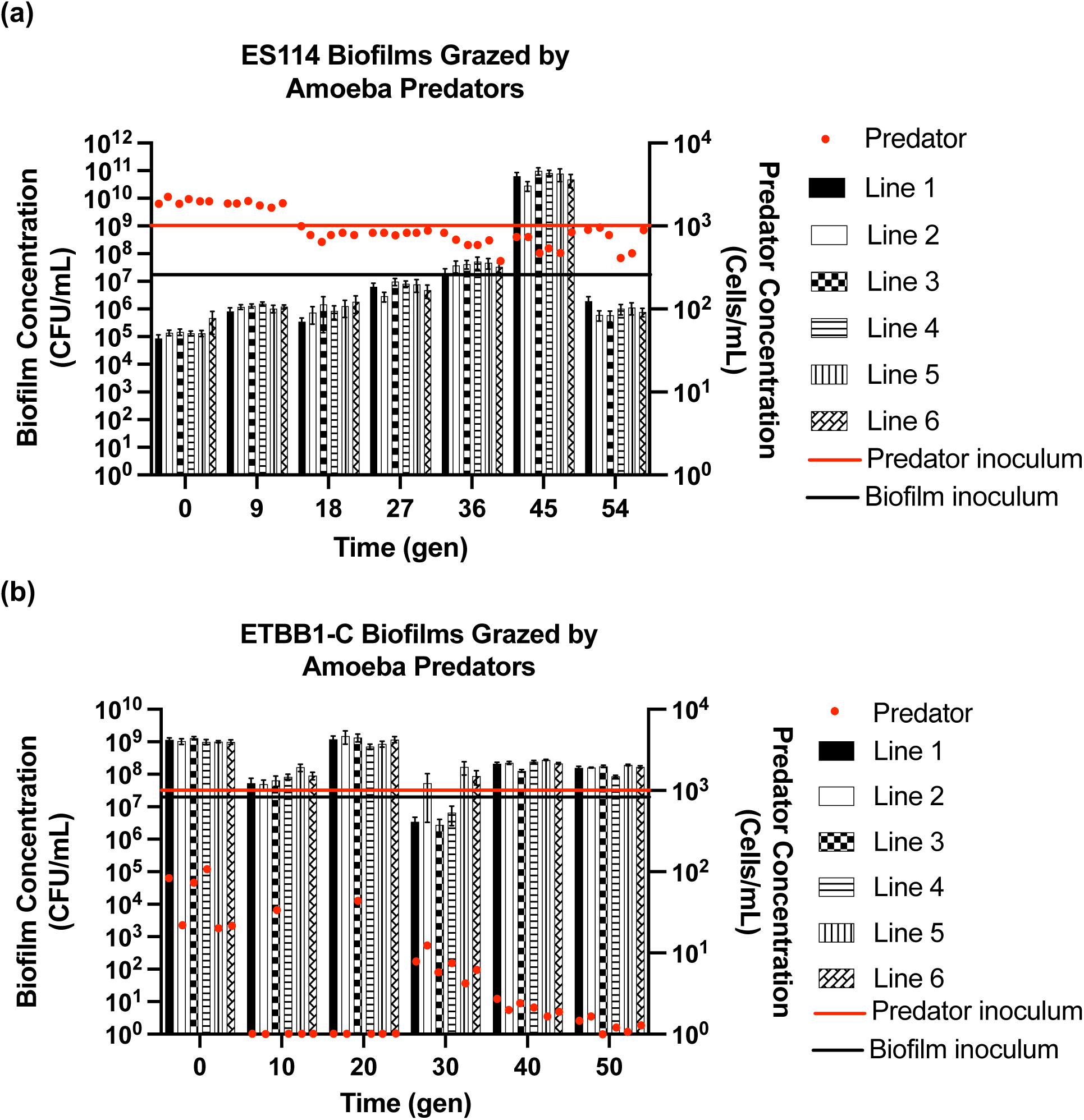
Concentrations of biofilm and amoeba populations measured in CFU/mL and cells/mL, respectively. Biofilm concentrations for six independent lines are measured against time in generations. The red line represents the initial 1,000 cells/mL inoculum of amoebas. The black line represents the initial 3 x 10^7^ CFU/mL biofilm inoculum. The red dot signifies the concentration of amoebas 24 hours after grazing on the biofilms. Each bar represents biofilm concentration of each line post-grazing. a) ES114 biofilms grazed by amoebas, b) ETBB1-C biofilms grazed by amoebas. Error bars are based on SEM with six biological replicates per lineage.

### Biofilm Assays

Six lineages of nongrazed ES114 biofilms were evolved under no predation selection up to 54 generations. The concentrations ranged from approximately 1 x 10^5^ to 1 x 10^8^ CFU/mL (Fig. 3a). The most significant change in biofilm concentrations occurs at generation 36. Lineage 6 in particular at generation 36 grew significantly compared to its ancestral counterpart with a p-value less than 0.0001 (Table 2). Likewise, six lineages of nongrazed ETBB1-C were evolved under no predation selection up to 50 generations. Biofilm concentrations varied less, ranging from around 1 x 10^7^ to 1 x 10^8^ CFU/mL (Fig 3b). The greatest difference in concentrations from the ancestors is observed in lineage 3 at generation 30 with a p-value of 0.0061 (Table 2).

**Table 2.** Symbols for Significance Levels.

| Symbol | Value |
| --- | --- |
| ns | $P > 0.05$ |
| * | $P \leq 0.05$ |
| ** | $P \leq 0.01$ |
| *** | $P \leq 0.001$ |
| **** | $P \leq 0.0001$ |

*V. fischeri* ES114 biofilms were grazed by *A. castellanii* up to 54 generations (Fig. 4a). ETBB1-C biofilms were grazed by *A. castellanii* up to 50 generations (Fig. 4b). ES114 biofilms were grazed by *T. pyriformis* up to 63 generations (Fig. 5a). *V. fischeri* ETBB1-C biofilms were grazed by *A. castellanii* and *T. pyriformis* up to 50 generations (Fig. 5b). Across all six lines, ES114 biofilms grazed by amoebas increase significantly in biofilm concentrations over time (Fig. 4a). For example, the concentration of biofilms for line 1 at 45 generations is significantly different from its concentration at 0 generations with a p-value less than 0.0001 (Table 3). However, they decrease drastically at the final measurement at generation 54. In contrast, amoebas appear to increase in population size compared to the initial inoculum. Across generations of grazing however, the amoeba populations decrease.

**Figure 5.**
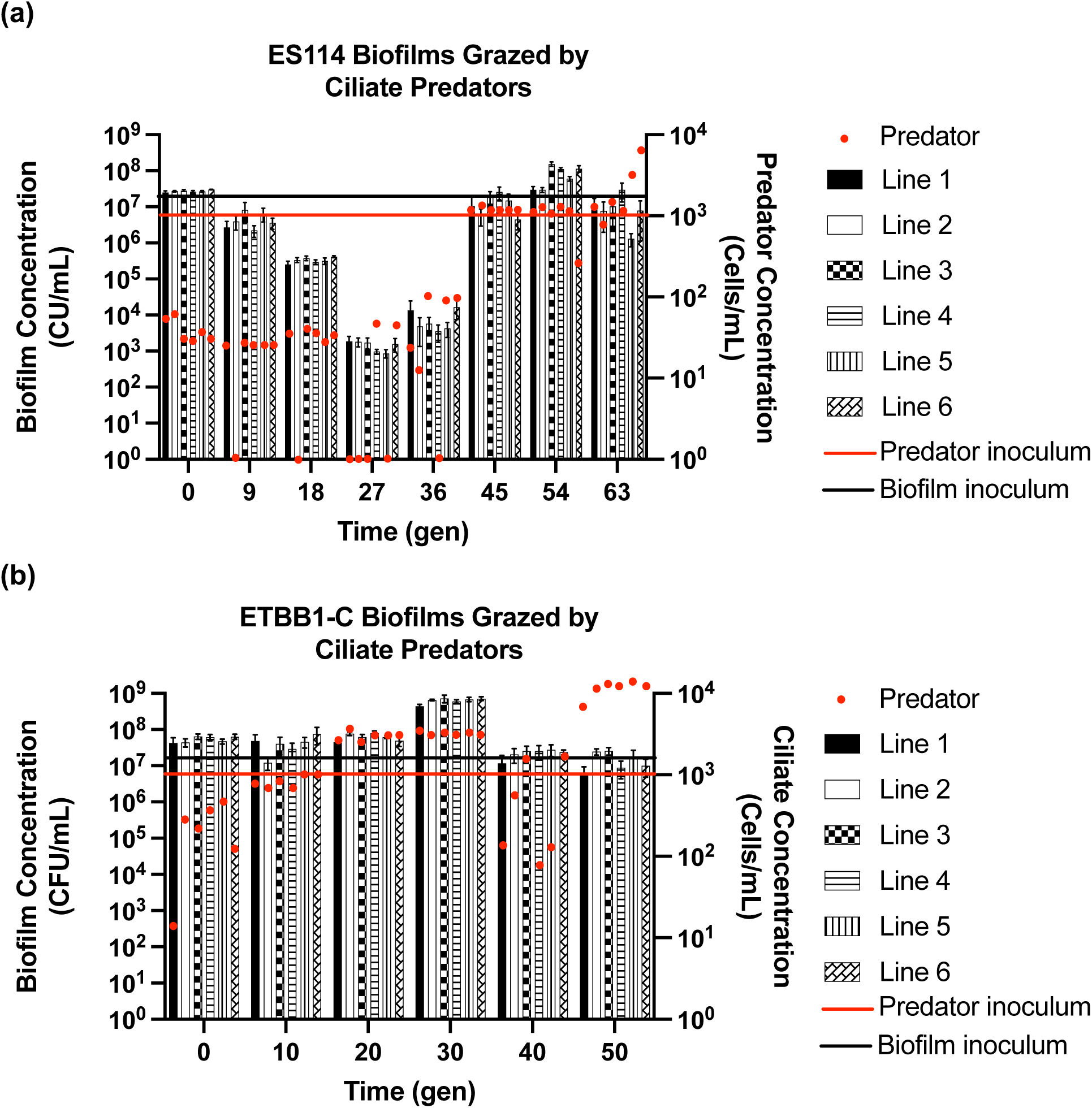
Concentrations of biofilm and ciliate populations measured in CFU/mL and cells/mL, respectively. Biofilm concentrations for six independent lines are measured against time in generations. The red line represents the initial 1,000 cells/mL inoculum of ciliates. The black line represents the initial 3 x 10^7^ CFU/mL biofilm inoculum. The red dot signifies the concentration of ciliates 24 hours after grazing on the biofilms. Each bar represents biofilm concentration of each line post-grazing. a) ES114 biofilms grazed by ciliates, b) ETBB1-C biofilms grazed by ciliates. Error bars are based on SEM with six biological replicates per lineage.

**Table 3.** P-values of Significance for Nongrazed Biofilm Concentrations.

| Comparison | Significance | P-value |
| --- | --- | --- |
| ES114 L6: Gn 36 vs. Ancestor | **** | <0.0001 |
| ES114 L6: Gn 54 vs. Ancestor | ns | 0.9827 |
| ETBB1-C L3: Gn 30 vs. Ancestor | ** | 0.0061 |
| ETBB1-C L3: Gn 40 vs. Ancestor | **** | <0.0001 |
| ETBB1-C L6: Gn 50 vs. Ancestor | * | 0.0361 |
| L6 Ancestor: ES114 vs. ETBB1-C | ns | 0.1553 |
| L6: ES114 Gn 36 vs. ETBB1-C Gn 30 | ns | 0.3313 |
| L6: ES114 Gn 54 vs. ETBB1-C Gn 50 | ** | 0.0017 |

### Bioluminescence Assays

The most evolved nongrazed ES114 lineages at generation 54 were brightest compared to its ancestor. The brightest lineage was line 1 with a p-value less than 0.0001 compared to the ancestor (Table 4). The brightest nongrazed ETBB1-C lineages were at generation 40, with the brightest one being from line 3 (Fig. 6b) with a p-value less than 0.0001 compared to the ancestor (Table 4). Bioluminescence for this strain appeared to decrease at the latest generation at generation 54. Nongrazed ES114 biofilms in general were less bright than their ETBB1-C counterparts.

**Table 4.** P-values of Significance for Grazed Biofilm Concentrations.

| **Comparison** | **Significance** | **P-value** |
| --- | --- | --- |
| ES114-AC G L1: Gn 45 vs. Ancestor | **** | <0.0001 |
| ES114-AC G L1: Gn 54 vs. Ancestor | ns | 0.9999 |
| ETBB1-C-AC G L3: Gn 30 vs. Ancestor | **** | <0.0001 |
| ES114-TP G L4: Gn 27 vs. Ancestor | ** | 0.0048 |
| ETBB1-C-TP G L3: Gn 30 vs. Ancestor | **** | <0.0001 |
| ES114 G L5 Gn 54: AC vs. TP | **** | <0.0001 |
| ETBB1-C G L4 Gn 50: AC vs. TP | **** | <0.0001 |
| AC G L4: ES114 Gn 54 vs. ETBB1-C Gn 50 | **** | <0.0001 |
| TP G L1: ES114 Gn 54 vs. ETBB1-C Gn 50 | ** | 0.0052 |

**Figure 6.**
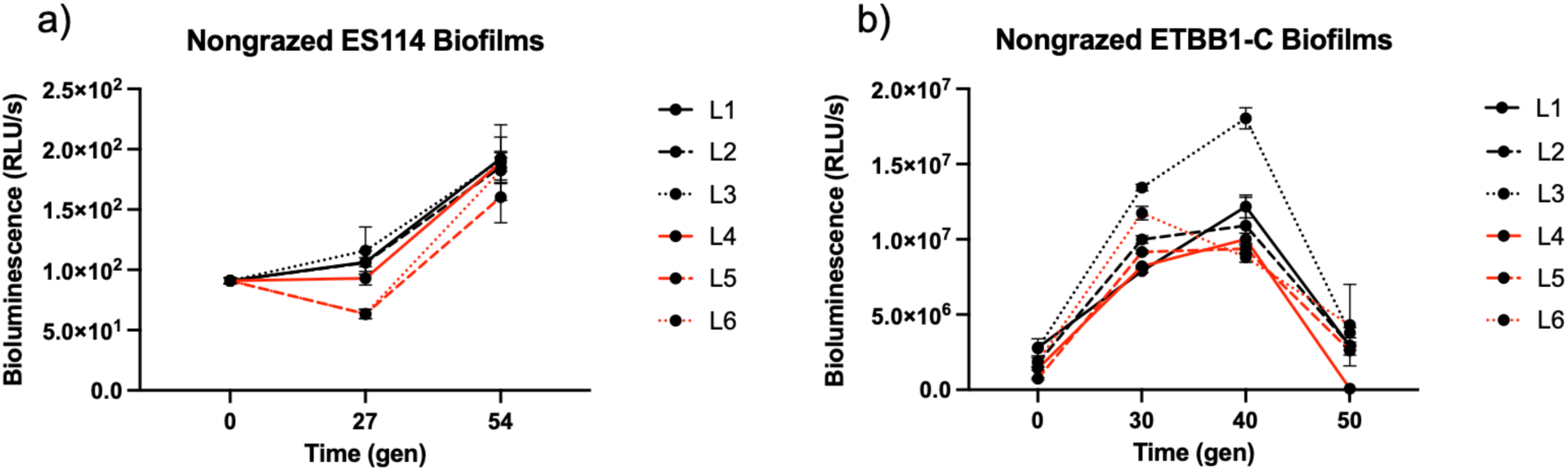
Bioluminescence assays of nongrazed a) ES114 and b) ETBB1-C biofilms for six independent lineages across generations. Error bars are based on SEM with three biological replicates.

ES114 biofilms when grazed by either amoebas or ciliates showed an increase in bioluminescence at the latest generation (Figs. 7a-b) with a p-value less than 0.0001 when compared to the ancestor for line 1 (Table 5). Grazed ETBB1-C biofilms overall were brightest at generation 40 (Figs. 7c-d). Line 6 of ETBB1-C biofilms grazed by amoebas was the brightest at generation 40 compared to the ancestors with a p-value less than 0.0001 (Table 5). However, for ETBB1-C biofilms grazed by ciliates, the brightest lineage was line 1 at generation 54.

**Figure 7.**
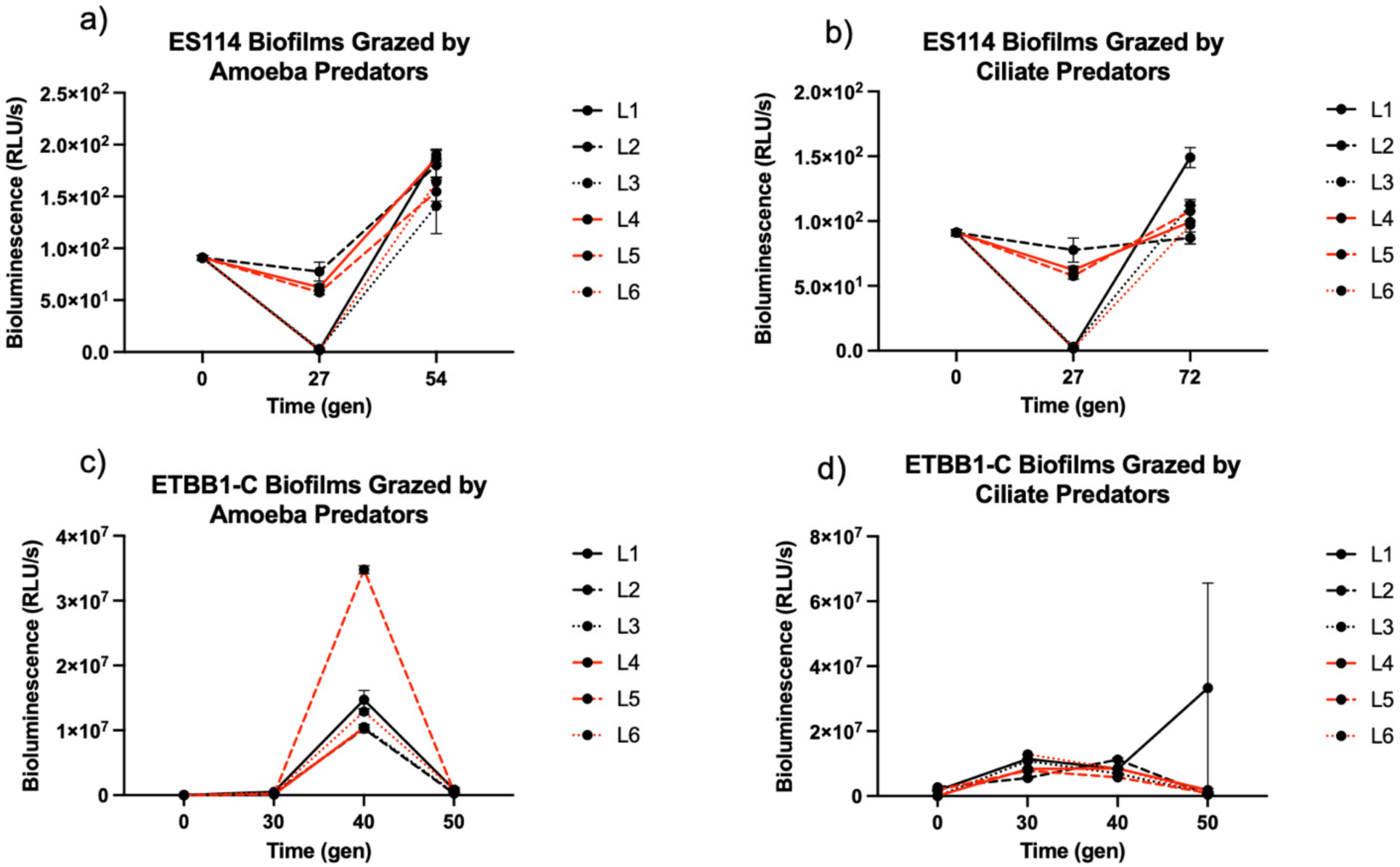
Bioluminescence assays of evolved *Vibrio fischeri* biofilms of a) ES114 grazed by *A. castellanii* and b) ES114 grazed by *T. pyriformis*, c) ETBB1-C grazed by *A. castellanii*, and d) ETBB1-C grazed by *T. pyriformis* for six independent lineages across generations. Error bars are based on SEM with three biological replicates.

**Table 5.** P-values of Significance for Nongrazed Biofilms Bioluminescence.

| **Comparison** | **Significance** | **P-value** |
| --- | --- | --- |
| ES114 L1: Gn 54 vs. Ancestor | **** | <0.0001 |
| ETBB1-C L3: Gn 40 vs. Ancestor | **** | <0.0001 |
| L3: ES114 Gn 54 vs. ETBB1-C Gn 50 | **** | <0.0001 |

**Table 6.** P-values of Significance for Grazed Biofilm Bioluminescence.

|  |  |  |
| --- | --- | --- |
| ES114-AC G L1: Gn 54 vs. Ancestor | **** | <0.0001 |
| ES114-TP G L1: Gn 72 vs. Ancestor | * | 0.012 |
| ETBB1-C-AC G L6: Gn 40 vs. Ancestor | **** | <0.0001 |
| ETBB1-C-TP G L1: Gn 40 vs. Ancestor | *** | 0.0006 |
| AC G L1: ES114 Gn 54 vs. ETBB1-C Gn 50 | * | 0.0165 |
| TP G L1: ES114 Gn 72 vs. ETBB1-C Gn 50 | ns | 0.4112 |

### Competition Indices

The competitive index between amoeba-grazed and nongrazed *V. fischeri* ES114 biofilms at lineage 1 for generation 54 was 5.55 (Fig. 8). Even though amoeba-grazed ES114 biofilms were lower at generation 54 compared to generation 45 (Fig. 4a), they were still greater in concentration compared to their nongrazed counterparts. Amoeba-grazed ES114 biofilms for lineage 1 at generation 54 was at 1.92 x 10^6^ CFU/mL (Fig. 4a). Nongrazed ES114 biofilms for this same lineage and generation was at 3.7 x 10^4^ CFU/mL. This difference was significant with a p-value less than 0.0001. Bioluminescence between these two ES114 strains however were not significantly different from each other.

**Figure 8.**
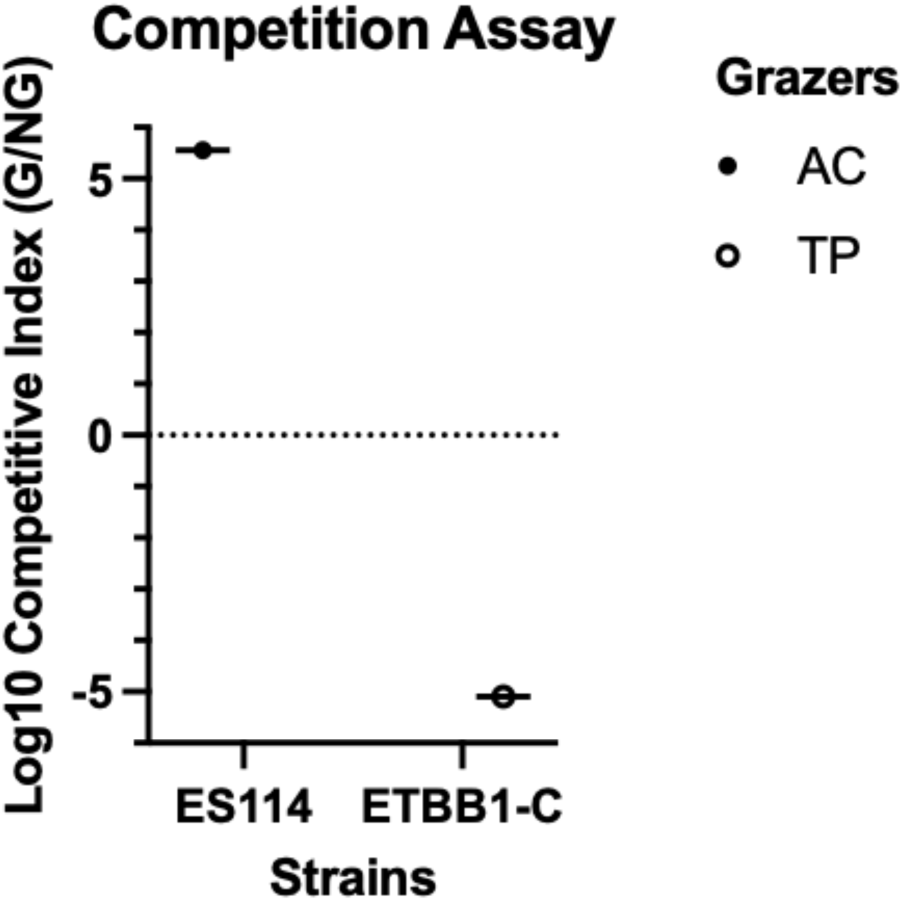
Competition assay showing two different competitive indices. One showing the competitive index between ES114 nongrazed and amoeba-grazed strains for line 1 at generation 54. The other competitive index is between ETBB1-C nongrazed and ciliate-grazed strain for line 2 at generation 50.

For *V. fischeri* ETBB1-C between ciliate-grazed and nongrazed at lineage 2 for generation 50, it was -5.097 (Fig. 8). It should be noted that the ciliate-grazed strain had a much lower biofilm concentration of 2.4 x 10^7^ CFU/mL in the grazing assay (Fig. 5b) compared to its nongrazed counterpart which had a biofilm concentration of 1.1 x 10^8^ CFU/mL (Fig. 3b). The difference was significant with a p-value less than 0.0001. In addition, the ciliate-grazed strain had lower bioluminescence with a value of 4.6 x 10^6^ RLU/s (Fig. 7d) compared to its nongrazed counterpart which had a bioluminescence value of 2.9 x 10^7^ RLU/s. This was significant with a p-value less than 0.0001.

## Discussion

### V. fischeri ES114 Biofilms show a cyclic response to predation

*V. fischeri* ES114 biofilms evolved under no predation stress exhibits a relatively stable population across generations (Fig. 3a). However, these biofilms show a cyclic response to predation by both *A. castellanii* and *T. pyriformis* (Figs. 4a and 5a). When *V. fischeri* ES114 biofilms are treated with *A. castellanii*, biofilm concentrations increase across generations. However, biofilm concentrations at evolved generations of grazing eventually begin to decrease. Grazing by *T. pyriformis* also exhibits a cyclic response albeit in the opposite manner. Biofilm concentrations decrease initially across generations, only then to increase up to the same levels as the ancestors, beginning at generation 36 (Fig. 5a). This cyclic response to predation is typical of predator-prey interactions (20), where prey (biofilms) responds to the constant pressure of grazing by either growing faster or releasing anti-grazing compounds (13, 21). The caveat to this behavior is that both populations are fluctuating in response to each other according to the Lotka-Volterra model. Our results from this experiment however do not show that as even though the biofilms appear cyclic, the protozoans either decrease or increase altogether across generations. This suggests that the biofilms may have a multifaceted response to grazing.

*V. fischeri* ES114 biofilms treated by *A. castellanii* do appear to follow the Lotka-Volterra model when biofilm concentrations are low and amoeba concentrations are high. Across generations, both populations show an inverse response to each other where the biofilm concentrations increased, but amoeba concentrations decreased. However, it should be noted that this phenomenon only occurs in biofilms that are being evolved. Protozoans used during each grazing experiment are from a stock culture (so these are not under experimental evolution selection) and the initial concentration remains the same during each grazing challenge. Grazing by *T. pyriformis* shows greater deviation from the Lotka-Volterra model in that both populations appear to change in size in the same direction as each other (Fig. 5a).

Differences in the cyclical response of *V. fischeri* ES114 biofilms to *A. castellenii* versus *T. pyriformis* may be due to the type of grazer. Some protozoans are known to induce biofilm formation depending on the age of the biofilm and the thickness of the community (22). For example, *A. castellanii* was shown to induce *Salmonella enterica* to form denser biofilms than their nongrazed counterparts under nutrient-rich conditions (23). It is plausible that *V. fischeri* ES114 biofilms are responding to amoeba-grazing by forming thicker biofilms. The decrease in biofilm concentrations however in the later generations may be a result of sloughing off of excess cells. This can occur when large particles of biofilm are released that are comparable to or greater than the thickness of the original biofilm (24).

However, *T. pyriformis* causes *V. fischeri* ES114 biofilms to fluctuate in cell density across generations cyclically as compared to grazing by *A. castellani*, which is more stable (Fig. 5a). *T. pyriformis*, as with other protists, have been shown to graze on bacterial biofilms (25). ES114 biofilms in previous grazing studies were also shown to be susceptible to *T. pyriformis* in contrast to free-living strains (13). However, *T. pyriformis* can also induce biofilm formation during grazing (26). The decrease in biofilm concentrations across early generations may show the susceptibility to grazing by *T. pyriformis*. In later generations, the increase in biofilm concentrations could be from an adaptive response to *T. pyriformis* predation, allowing the biofilm to stabilize or grow back faster than the initial generation.

There have been multiple observations for various biofilms being induced by *T. pyriformis*. Confocal laser scanning microscopy has shown *T. pyriformis* cells on the surface of *B. cenocepacia* biofilms or embedded in their matrix influences the concentration and thickness during grazing experiments (27). Embedded *T. pyriformis* was found to have *B. cenocepacia* cells on them as well as inside of their food vacuoles, which are known to withstand digestion. These ciliates are potentially lysed by biofilm-produced toxins, and release their internal content (including the intact *B. cenocepacia*), which are thought to act as nutrients to further aid in biofilm formation (27). Therefore, it was proposed that *T. pyriformis* cells acted as centers of new biofilm being formed or regenerated. This may be the case of *V. fischeri* ES114 biofilms where they adapted to grazing pressure by forming biofilms on these ciliated predators while potentially embedded in the biofilms. Previous studies have observed that *Tetrahymena* cells were killed after grazing and eventually absorbed by *V. fischeri* biofilms (13). This “fight back” behavior would allow bacteria such as *V. fischeri* to survive and possibly gain additional carbon and nitrogen sources by the dead protists.

### Resilience Prevents Grazing on V. fischeri ETBB1-C Biofilms

In contrast to *V. fischeri* ES114, biofilms produced by *V. fischeri* ETBB1-C are generally greater in concentration and more stable across generations (Fig. 3b). Nongrazed *V. fischeri* ETBB1-C biofilms appear to remain around 5 x 10^7^ CFU/mL. Protozoan grazing appears to perturb biofilm populations slightly. Across generations, *A. castellanii* in general appears to induce *V. fischeri* ETBB1-C biofilm formation in comparison to their nongrazed counterparts. The amoeba predator populations on the other hand decrease significantly from their initial inoculum (Fig. 4b).

Reduction in amoebas post-grazing on *V. fischeri* ETBB1-C biofilms may be a result of exposure to toxins produced by the biofilms. One such toxin that is known to be produced by biofilms is violacein that can act as anti-predatory measures (13, 28). In the closely related species *V. cholera*, Lipid A of the lipopolysaccharide (LPS) is thought to inhibit phagocytosis by *T. pyriformis* (29). LPS itself is also found to protect *Klebsiella pneumoniae* from the slime mold, *Dictyostelium discoideum* (30). *V. fischeri*, as with most Gram-negative bacteria, has lipid A as part of the LPS outer membrane layer (31). Additionally, it has been found that the outer membrane protein OmpU in *V. cholerae* has a role in resistance against digestion by protozoans (32). OmpU is thought to elicit a sigma factor σ^E^ mediated stress response when it interacts with host antimicrobial peptides. *ompU* is also found in *V. fischeri* and has a significant role in squid colonization (33). This gene codes for OmpU, an adhesion protein that acts as a virulence factor for *Vibrio* species in colonizing their host (34). It is thought that OmpU for *V. fischeri* is needed for adherence in colonizing their squid host. So, it is likely that *V. fischeri* also uses these molecules as a defense mechanism against protozoan predation. Future studies examining the expression patterns of these genes will uncover which mechanisms are under selection for grazing by these predators.

Grazing by *T. pyriformis* had little impact on ETBB1-C biofilm formation except for at generation 30 (Fig. 5b). Regardless, their populations are not as reduced compared to populations of *A. castellanii* when grazing on ETBB1-C biofilms. *T. pyriformis* are typically suspension-feeding protists (35) and swim through the suspension eating cells that are in their free-living state. Therefore, it is likely that *T. pyriformis* predators are feeding on planktonic cells being sloughed off from biofilms rather than from the biofilms themselves. For example, *T. pyriformis* was found to graze upon planktonic cells of *Burkholderia cenocepacia*, while at the same time stimulating biofilm formation (27). Similarly, *Xanthomonas retroflexus* and *Stenotrophomonas rhizophila* increased in biofilm formation when treated with *T. pyriformis* (36), even though their planktonic cells were grazed by *T. pyriformis*. *Paramecium tetraurelia*, a ciliated protist akin to *T. pyriformis*, was also found to induce biofilm formation in *Pseudomonas putida*, yet grazed upon their planktonic cells (26).

### Comparative analysis between V. fischeri ES114 and ETBB1-C biofilm response to grazing

Protozoan predation is known to affect biofilm formation in many other systems (21). In fact, it has been found to induce virulence in pathogenic bacteria (37). Previous studies demonstrated that *V. cholerae* grazed by *A. castellanii* upregulate virulence genes, making them more pathogenic (37). Whether grazing selects for symbiotic microbes to be more beneficial is illuminated from this study. For instance, the results from this study show certain strains are more resilient to grazing than others.

*V. fischeri* ETBB1-C produces biofilms resilient to both types of predators. This is further supported by the fact that the amoebas when grazing on ES114 biofilms remained close to the initial inoculum post-grazing whereas when grazing on ETB11-C biofilms, they decreased significantly in concentration. This observation in part may be explained by differences in biofilm morphology between ES114 and ETBB1-C. *V. fischeri* strains isolated from *E. tasmanica* (ETBB1-C) have been shown to grow denser biofilms than *E. scolopes* strains (6, 13). However, when evolved by passaging through *E. tasmanica* squids, Hawaiian *V. fischeri* ES114 becomes better at forming biofilm (6). By forming thicker biofilms, the amoebas might lack access to the cells within the biofilm (38), and therefore only graze the surface of thicker biofilms. Similarly, the biofilms may increase its production of extracellular polysaccharide (EPS) to shield itself from predation (22). The combination of forming denser biofilms in addition to increased production of EPS in response to grazing can potentially create a shield preventing protists from grazing (38). The decrease in protozoan cells as demonstrated in our study therefore may be a result of starvation or from being neutralized by toxins produced by biofilms released upon grazing (30). Ciliated protists such as *T. pyriformis* are thought to induce biofilm formation from a number of biofilm grazing studies through different means ranging from recycling nutrients in the liquid medium by the movement of their cilia (36), to acting as a source of nutrients themselves when lysed (27).

Morphological changes also occur, such as a shift from smooth to wrinkled biofilms, protecting biofilms from surface-feeding protists such as *Rhynchomonas nasuta* and *A. castellanii.* This morphological difference may cause protozoans to starve due to inability to access and digest parts of the biofilm. In *V. fischeri*, the EPS production is regulated by the *syp* locus (39). The sigma factor σ^54^, a regulator of the *syp* locus, is known to control EPS production (40), and may regulate the way biofilms respond not only to the daily release of free-living cells, but also to the density of components that comprise the biofilm. The *sypD* gene contains the EpsG domain, which is homologous to the EpsG protein responsible for EPS production in *Methylobacillus* sp. (41). These genes may be involved in producing EPS, which helps protect biofilms from being grazed by protozoans. This can be attributed to the cell-linking fibrillar matrix produced by the EPS, potentially shielding biofilms from a wide variety of protists (42).

Other potential responses of ciliate-induced biofilm formation are the utility as an active defense mechanism to predation (36). For instance, the wrinkly biofilms of *V. cholerae* were selected in response to grazing through production of exopolymer (38). This phenotype along with anti-protozoan compounds contributed to *V. cholerae* biofilm resistance to grazing. Ciliate grazing can also stimulate a passive mechanism whereby the ciliary movement of this type of protozoan predator creates a current in the environment, increasing the movement of nutrients near the biofilm. This behavior increases the availability of nutrients to the biofilm surface, and may help stimulate their growth (26, 36). Such mechanisms in combination with other biofilm responses can provide multiple aspects of stimulation for biofilm formation, which in the natural environment can always be changing due to abiotic factors effecting both predator and biofilms.

Another abiotic factor that is known to impact these biofilm-predator interactions is temperature. Biofilms generally colonize a surface earlier and grow thicker under warmer temperatures (43). However, it is also shown that grazing by protozoans increases with elevated temperatures. This is explained by warming temperatures accelerating colonization by grazing protists, via increased growth rates and metabolism (43, 44). This results in changes in population dynamics between algae, bacterial biofilms, and grazing protists. Increasing temperatures promotes the photosynthetic rates of algae, causing them to release exudates which are then taken up by bacterial biofilms as nutrients (43). This causes biofilms to increase production of EPS which itself promotes growth of protists, in particular those belonging to the subclass peritrichia. These protists are filter-feeding ciliates which generally feed on bacteria in the water column, but have been found to live inside biofilm matrices via confocal laser scanning microscopy (43). Strong currents produced through the beating of their cilia may be enough to dislodge biofilm cells at warmer temperatures, allowing them to feed on these sloughed-off cells. This effectively leads to both a reduction in biofilm structures and increased grazing on the dislodged planktonic cells (43).

### Strain-Dependent Effects on Bioluminescence

Strain type appeared to have a bigger impact on bioluminescence than the type of grazer that were examined on all biofilms. *V. fischeri* ES114 increased in bioluminescence across generations under no protozoan treatment (control, Fig. 6a) as well as when treated with either *A. castellanii* or *T. pyriformis* (Figs. 7a-b). However, in both cases, bioluminescence decreased in three lineages at generation 27. This may be due to a reduction in biofilm density at this generation at least in the case for *T. pyriformis* grazed biofilms (Fig. 5a). As for *A. castellanii* grazed *V. fischeri* ES114 biofilms, there is a possibility that some of the biofilm lineages decreased in bioluminescence perhaps as a stress response to predation. It is known that other stressors such as toxins reduce bioluminescence in *V. fischeri* (45), and further examining of the expression of such toxins will enlighten our understanding of this phenotype.

### Grazer-Dependent Effects on Host Colonization

Predation also has an impact on host colonization. *V. fischeri* can form biofilms outside of the squid host and subsequently be affected by protozoan predation (13). Just how susceptible *V. fischeri* biofilms are to protozoan predation appears to be dependent on a multitude of factors. For example, the effects of protozoan predation can differ between specific strains of *V. fischeri.* The Hawaiian isolate ES114 is more susceptible to predation, showing drastic changes in biofilm concentration in response to both protozoans, *A. castellanii* and *T. pyriformis*. *A. castellanii* has an insignificant effect on bioluminescence, but a significant effect on colonization. Amoeba-grazed ES114 biofilms for lineage 1 at generation 54 were more competitive in colonizing their squid host compared to their nongrazed counterparts (Fig. 8). This may be related to the evolved biofilms being able to form denser biofilms than their nongrazed counterparts. During the colonization process, the amoeba-grazed strains may have been able to form thicker aggregates around the pores of the light organ, thereby outcompeting their less dense nongrazed counterparts. It is well known from previous studies that strains that cannot form aggregates fail to colonize the squid host compared to those that can form aggregates outside of the pores (46). Previous work has shown that *V. fischeri* can be evolved to become better colonizers to their squid host (6, 47). This work also demonstrates that protozoan predation by amoebas can evolve *V. fischeri* to become better colonizers to their squid host.

Protozoan predation however could also negatively impact colonization fitness. For instance, it was found that ciliate predation dampens the colonization fitness of ETBB1-C. Ciliate-grazed ETBB1-C biofilms for lineage 2 at generation 50 were less competitive than their nongrazed counterparts in colonizing their squid host (Fig. 8). This may be in due to their loss in both biofilm forming and bioluminescence phenotypes as mentioned earlier. As with biofilm formation, bioluminescence also plays a role in colonization fitness. It is well known in previous studies that mutant *V. fischeri* strains deficient in bioluminescence termed as “dark mutants” were outcompeted by their brighter wildtype counterparts (48). Therefore, the loss in biofilm formation and bioluminescence caused by grazing by *T. pyriformis* may have contributed to the loss in colonization fitness.

## Conclusion

This study demonstrates the importance of how biotic factors such as protozoan predation can influence the regulation of an important symbiotic characteristic such as biofilm formation. In the squid-*Vibrio* beneficial association, biofilm-like aggregates are required for the squid host to be colonized and *Vibrio* bacteria to persist. For example, previous studies have demonstrated that biofilm formation and motility are inversely related, where bacteria that form dense biofilms tend to be poor in motility (6, 49, 50). The opposite is also true where poor biofilm-forming bacteria tend to be robust in motility. In addition, bioluminescence is directly proportional to cell density. That is, as *V. fischeri* populations become denser, so does bioluminescence (51). This is vital in the context of symbiosis between the squid and their *Vibrio* symbiont as both these of these characteristics are needed for proper colonization (52).

Overall, this work has shown that *V. fischeri* biofilms outside of the squid host are affected by protozoan predation, which subsequently has a fitness cost for colonizing and persisting in symbiosis with their squid host. Biotic stressors in the surrounding environment can act as one of many ecological factors that impact environmentally transmitted symbiosis. This delicate balance between fitness trade-offs when enduring selection pressures such as grazing outside the squid host is crucial to having a successful symbiosis, where colonization in the squid host is drastically affected by continual grazing in the free-living state.

Future plans are to examine biofilm regulatory loci to determine if indeed the expression of genes involved in biofilm formation and EPS production such as *sypD* are up-regulated in strains evolved in response to protozoan predation. Other key loci would include those involved in resistance against protozoan predation such as *lpxB* and *ompU*. Additionally, given the time-intensive nature of these grazing experiments, mathematical modeling beyond thousands of generations can provide additional clues on what factors are most important during long term experimental evolution grazing experiments. This would include minute details such as biofilm and protozoan grazing rates, both of which can be parameterized to predict the long-term outcome of such environmental selection pressures (26). Long term evolution in response to strong selection pressures before colonization can give us insight on how this may affect the overall fitness of the symbiosis between sepiolid squids and *V. fischeri*, and whether these factors contribute to the overall persistence of a successful beneficial association.

## Acknowledgements

The work was supported by NSF DBI-2214028, NASA EXO 80NSSC18K1053 and the School of Natural Sciences at UC Merced to M.K.N. N. Mora, H. Owen and A. Shakra assisted with biofilm assays and statistical analyses.

